# The Genetic Architecture of Leaf Stable Carbon Isotope Composition in *Zea mays* and the Effect of Transpiration Efficiency on Elemental Accumulation

**DOI:** 10.1101/2020.03.12.989509

**Authors:** Crystal A. Sorgini, Lucas M. Roberts, Asaph B. Cousins, Ivan Baxter, Anthony J. Studer

## Abstract

With increased demand on freshwater resources for agriculture, it is imperative that more water-use efficient crops are developed. Leaf stable carbon isotope composition, δ^13^C, is a proxy for transpiration efficiency and a possible tool for breeders, but the underlying mechanisms effecting δ^13^C in C_4_ plants are not known. It has been suggested that differences in specific leaf area, which potentially reflects variation in internal CO_2_ diffusion, can impact leaf δ^13^C. However, at this point the relationship has not been tested in maize. Furthermore, although it is known that water movement is important for elemental uptake, it is not clear how manipulation of transpiration for increased water-use efficiency may impact nutrient accumulation. Here we characterize the underlying genetic architecture of leaf δ^13^C and test its relationship to specific leaf area and the ionome in four biparental populations of maize. Five significant QTL for leaf δ^13^C were identified, including both novel QTL as well as some that were identified previously in maize kernels. One of the QTL regions contains an Erecta-like gene, the ortholog of which has been shown to regulate transpiration efficiency and leaf δ^13^C in *Arabidopsis*. Our data does not support a relationship between δ^13^C and specific leaf area, and of the 19 elements analyzed, only a weak correlation between molybdenum and δ^13^C was detected. Together these data begin to build a genetic understanding of leaf δ^13^C in maize and suggest the potential to improve plant water use without significantly influencing elemental homeostasis.

**Article Summary:** Quantitative genetics approaches were used to investigate the genetic architecture of leaf stable carbon isotope discrimination (δ^13^C) in maize. Developing a better understanding of leaf δ^13^C could facilitate its use in breeding for reduced transpirational water loss. Several genomic regions were identified that contribute to the variation observed in leaf δ^13^C. Furthermore, contrary to what has been observed in other species, leaf δ^13^C was not correlated with specific leaf area. Finally, a leaf ionomic analysis indicates that a reduction in transpiration, and thus mass flow, would not result in a decrease in nutrient accumulation.

## INTRODUCTION

The impacts of global population growth and climate change on natural resources indicate that the future of food security will depend on increasing both the productivity and sustainability of agriculture systems (National Academies of Sciences, Engineering, and Medicine, 2018). Improving crop water-use efficiency (WUE) would ameliorate the effects of the increasing frequency and severity of droughts (Scheffield and Wood 2008; Chapman *et al.* 2012, Leakey 2019). Agronomic WUE can be defined as the amount of yield, whether grain or biomass, produced per the total amount of water utilized by the crop (Condon *et al.* 2004). Many factors can affect WUE including transpirational water loss through the stomatal pores on the leaf’s surface. In C_3_ plants the amount of carbon available for assimilation is limited by stomatal and mesophyll conductances to CO_2_ (Flexas *et al.* 2016) and therefore correlated to the rate of transpiration. For example, yield was shown to be positively associated with cumulative transpiration in soybean (Purcel 2007), and higher net carbon assimilation was accompanied by higher transpiration in rice (Adachi *et al.* 2017). However, higher rates of biomass yield do not always correspond to higher transpiration rates in C_4_ plants due to the evolution of the carbon concentrating mechanism. The uncoupling of CO_2_ assimilation and transpiration has been demonstrated in field and greenhouse grown maize (Walker 1986, Kolbe *et al.* 2018a). Thus, there is the potential to increase transpiration efficiency, or carbon gain per amount of water transpired, without reducing productivity in C_4_ species (Leakey, 2019). A large amount of variation is present in the transpiration rates of C_4_ crop species, including sorghum (Hammer *et al.* 1997) and maize (Bunce 2010), suggesting that existing occurring alleles could be exploited for optimizing WUE.

Although increasing transpiration efficiency provides a strategy to avoid the negative effects of water limitation on plant growth and development (Passioura 1996, Chaves *et al.* 2002, Jaleel *et al.* 2009), there is the possibility of pleiotropic side effects given the fundamental requirement for water movement in plants. A potential impact of reducing transpiration could be a corresponding reduction in the uptake and mobilization of water-soluble nutrients. As water is absorbed by roots, nutrients in solution come in contact with the root surface in a process known as mass flow (Barber *et al.* 1963). Most nutrients are acquired by mass flow, although phosphorus is a notable exception that contacts the root through diffusion (Barber *et al.* 1962).

Reducing transpiration may also affect nutrient uptake facilitated by symbiosis with mycorrhizal fungi (Marschner and Dell 1994). Therefore, the manipulation of basic plant processes such as transpiration for improved WUE must also consider potential impacts on the availability of essential plant nutrients. Previous research has shown in the C_4_ plant sorghum that total leaf mineral content is positively correlated to transpiration efficiency (Masle *et al.* 1992). While meta-analyses of high CO_2_ grown plants with reduced transpiration have shown a drastic reduction in nutrient accumulation in C_3_ crops (McGrath and Lobell 2013), sorghum showed no difference and maize had similar levels of zinc, protein, and phytate, but a decrease in iron accumulation (Meyers *et al.* 2014). Although part of the reduction in nutrient content can be explained by dilution, due to increased growth at high CO_2_, this does not completely account for the observed reduction. An ionomics (high-throughput elemental profiling) approach has been used in maize to assess kernel nutrient content (Baxter *et al.* 2014). A similar ionomics approach in leaf tissue could be used to assess the effect of transpiration on nutrient uptake.

The difficulty and labor-intensive nature of accurately quantifying the amount of water that an individual plant transpires has been a major limitation to breeding for transpiration efficiency. This has resulted in the selection for drought tolerance rather than applying a direct selection for water use. One alternative method is the use of leaf stable carbon isotopes as a proxy for transpiration efficiency. The stable carbon isotope composition, δ^13^C, reflects the amount of ^13^C present in plant tissue relative to a standard (Keeling 1979). Enzymes in the process of carbon fixation discriminate differently against the heavier ^13^C atoms in a process known as fractionation (Farquhar *et al.* 1982, O’Leary 1988). It has been widely shown that stable carbon isotopes can be used as a proxy trait for quantifying a plant’s transpiration efficiency in C_3_ plants (Farquhar *et al.* 1989a, Farquhar *et al.* 1989b, Condon *et al.* 1990, Virgona *et al.* 1990, Condon *et al.* 1993, Barbour *et al.* 2010) and in C_4_ plants (Henderson *et al.* 1998, von Cammerer *et al.* 2014, Ellsworth *et al.* 2017, Twohey III *et al.* 2019, Ellsworth *et al.* 2020). Studies have also shown that δ^13^C can be influenced by environmental factors including light intensity and drought (reviewed in Cernusak *et al.* 2013). However, the genetic control of δ^13^C remains unknown in C_4_ species.

Kolbe *et al.* showed that δ^13^C did not correlate with any of the photosynthetic enzymes previously posited to control for δ^13^C variation (2018b). Additionally, a transcriptome analysis was unable to identify a clear candidate gene (Kolbe *et al.* 2018b). Quantitative genetic approaches have the potential to reveal the genetic control of δ^13^C in C_4_ species because genomic locations are tested for associations with the trait of interest, without *a priori* knowledge of the mechanism underlying the variation. Maize is ideal for use in mapping studies due to its high level of recombination and low linkage disequilibrium (Yu and Buckler 2006). Mapping methods have been successfully used for decades to identify genes controlling complex traits in maize, with evolving approaches to tackle more difficult traits (Wallace *et al.* 2014). In addition, maize is both a model organism with available populations and genomic data, and one of the three most important global crops contributing to 30% of the total calories consumed by humans (Shiferaw *et al.* 2011).

There have been several previous studies that used quantitative genetics to investigate δ^13^C in C_3_ species (Teulat *et al.* 2002, Masle *et al.* 2005, Rebetzke *et al.* 2008, Xu *et al.* 2009). In Arabidopsis the gene *ERECTA* was identified in a QTL study for isotopic discrimination and was found to alter transpiration efficiency by altering stomatal density (Masle *et al.* 2005). Genetic mapping of leaf δ^13^C has also been performed in the C_4_ species *Setaria viridis* (Feldman *et al.* 2018, Ellsworth *et al.* 2020) and kernel δ^13^C has been mapped in the C_4_ maize (Gresset *et al.* 2014, Avramova *et al.* 2019). Although the QTL found for C_4_ species still require fine-mapping to identify the causative gene, no correlation was observed between kernel δ^13^C and leaf δ^13^C (Foley 2012). The lack of correlation may be the result of post-photosynthetic fractionation (Badeck *et al.* 2005), and therefore mapping QTL for δ^13^C in leaves may reveal additional loci not found using kernels. In this manuscript, we focus on leaf δ^13^C in maize and its association with leaf elemental composition. We also investigate variation in specific leaf area (SLA) and its potential relationship to leaf δ^13^C by CO_2_ diffusion. Characterization of the genetic architecture of leaf δ^13^C will provide a better context for understanding what drives δ^13^C, which will allow breeders to utilize this trait in crop improvement.

## MATERIALS AND METHODS

### Plant Material

All experiments were planted at the University of Illinois Crop Sciences Research Farm, Urbana IL and were subject to natural conditions without supplemental irrigation. NAM RIL families CML103, CML333, NC358, and Tx303 and NAM founder parents (McMullen *et al.* 2009) are publicly available through the Maize Genetic Cooperative Stock Center. The NAM RIL families were planted in the summer of 2015 using an augmented incomplete block design. For this experiment fifteen kernels were planted in each 3.7 meter row with 0.8 meter spacing between rows and 0.9 meter alleys. The families were randomized together, with each block consisting of 20 lines and 2 checks (B73 and one of the other founder lines). Of the 880 plots, 10% were dedicated to checks with the common parent B73 appearing in 40 plots and each of the four founder lines appears in ten plots. All lines used for the GWAS experiment are publicly available through the USDA Germplasm Resources Information Network (GRIN). This experiment was planted on May 23^rd^ 2016. Twenty kernels were planted in each 3.7 meter row with 0.8 meter spacing between rows and 0.9 meter alleys, and then thinned to 15 plants per row. A complete list of lines used can be found in Tables S1-S3.

### Tissue Sampling

Samples for δ^13^C analysis from the NAM RIL populations were collected six weeks after planting as follows. A rectangular piece of tissue approximately 7.5 cm X 5 cm was taken from the center of the leaf blade of the uppermost fully expanded leaf from four plants in each plot. Samples were placed in a coin envelope and dried at 65^°^C for at least 7 days. After drying four hole punches (each 0.058532 cm^2^) were taken and placed in a 6 mm x 4 mm tin capsules (OEA Laboratories # C11350.500P) for analysis using a Delta PlusXP (Washington State University) isotope ratio mass spectrometer. Leaf samples to measure specific leaf area (SLA) were collected from four plants in each of the plots (preferentially but not necessarily the same plants as were collected for δ^13^C) using a 1.6 cm diameter cork borer. Leaf discs were dried at 65^°^C for at least 7 days prior to weighing on an analytical balance (Model MS204S). Specific leaf area (SLA) was calculated as the area of a leaf disc divided by its dry weight. These same leaf discs were then used for ionomics analyses as described in Pauli *et al*. 2018. Leaf samples for δ^13^C analysis from the GWAS panel were collected from the uppermost fully expanded leaf seven weeks after planting using the hole punch method and processed as described previously (Twohey III *et al.* 2019). Due to the high level of diversity in this panel, some lines were flowering when samples were collected, which resulted in tissue being collected from the flag leaf. These samples were analyzed using a Costech instruments elemental combustion system and a Delta V Advantage isotope ratio mass spectrometer.

### Statistical Analysis

All analyses were completed using custom scripts and statistical packages in R (R Core Team 2017).

#### Correlation analysis

Pearson correlations using phenotype mean values were calculated with corr.test() in R package ‘psych’ (Revelle 2018) using complete observations and Holm’s method (Holm 1979) to adjust p-values for multiple testing. The correlation matrix was visualized using pairs.panel() in the R package ‘psych’ (Revelle 2018).

#### Stepwise regression QTL mapping

The analysis was completed using NAM_phasedImputed_1cM_AllZeaGBSv2.3 dataset. The file contains fully imputed and phased genotypes for most of the RILs in the NAM population (Zhao *et al.* 2006; Lipka *et al.* 2015). This HapMap format file was converted to numeric format where 0 is the B73 homozygote reference, 1 is a heterozygote, and 2 is the homozygote alternative parent. Phenotypic means were regressed onto genotype. Lowest p-values from the ANOVA values of the linear model were recorded (i.e. pvalues[i] = anova(lm(mypheno∼geno[i,])). The previously identified marker was added to the model and re-run in a stepwise regression procedure. Significance thresholds were determined by 200 permutations and alpha was set at 0.05. All analysis was completed using custom scripts in R (R Core Team 2017). Results were then compared to composite interval QTL mapping completed in R package ‘r/QTL’ (Broman *et al.* 2003).

#### Joint linkage mapping

The analysis was completed using HapMapv2 (Chia *et al.* 2012). The genotypic dataset consisted of 836 markers were scored on 624 RILs from four biparental families with B73 as a common parent. The marker subset represented markers that could be placed unambiguously on the physical map. Unambiguous markers are defined by those anchored in CDS positions of genes that have held consistent over genome versions verified by MaizeGDB cross reference tables. Missing data were imputed as previously described in Tain *et al.* (2011). Joint linkage models were constructed using custom script in R (R Core Team 2017) by a stepwise regression procedure. In general, we used linkage to test every marker across all four families to find the most significant QTL. The model has a family term and a marker:family term. The family term accounts for differences in mean phenotype between families. Inclusion of the marker:family term means that for each QTL we are assigning a separate effect to each family. The family term was included in the model and each of the 836 possible marker-by-family terms were assessed. Lowest p-values from the ANOVA values of the linear regression model were recorded (i.e. JL_pvalues[i] = anova(lm(my_pheno∼family+geno[,i]:family)). All 836 marker-by-family terms were tested. SNP effects were nested in families to reflect the potential for unique QTL allele effects within each family. The lowest resulting *p*-value was recorded for each permutation. Significance thresholds were determined using 1000 permutations for each family independently and alpha was set at 0.05.

#### Genome wide association stud

A subset of 413 of 503 diverse lines from Hirsch et. al 2014 that included the Wisconsin Diversity Set of Hansey et al 2011 was grown in 2016 and listed in Table S2. Hirsch et. al 2014 collected RNA from whole seedling tissue which was sequenced via IlluminaHiSeq and filtered to create a working set of 485,179 SNPs that is available at https://datadryad.org//resource/doi:10.5061/dryad.r73c5. The 413 lines were grown in 2016 and tissue was sampled when B73 was at the developmental stage V10. Isotopic analysis is described above. A genome wide association analysis was run using R package ‘GAPIT’ (Lipka *et al.* 2012) on leaf δ^13^C. Removal of SNPs with a minor allele frequency of less than 0.05 resulted in a subset of 438,222 SNPs being used in this analysis. A MLM model was used with model selection set to true to find the optimum number of principal components to account for population structure (Lipka et. al 2012). Significance threshold were calculated using the Bonferroni correction of familywise error rate. An alternative significance test was calculated using the Benjamini-Hochberg procedure for controlling the false discovery rate (Benjamini & Hochberg 1995).

### Data Availability

Genotypic datasets were downloaded from Panzea CyVerse iPlant Data Storage Commons (http://datacommons.cyverse.org/browse/iplant/home/shared/panzea). All phenotypic datasets were quality controlled for complete technical replicates, outliers, and availability of genotypic data. A list of all genotypes used in each analysis is provided in Tables S1-S4 and have been uploaded to figshare. Briefly, the δ^13^C analysis was completed with 640 RILs; including 156 CML103 RILs, 160 CML333 RILs, 159 NC358 RILs, and 165 Tx303 RILs (Table S1). The element analysis was completed using a total of 704 RILs; including 175 CML103 RILs, 181 CML333 RILs, 175 NC358 RILs, and 173 Tx303 RILs (Table S1). The SLA analysis used a total of 683 RILs; including 172 CML103 RILs, 176 CML333 RILs, 168 NC358 RILs, and 167 Tx303 RILs (Table S1). The Joint linkage analysis was completed using a total of 624 RILs; including 154 CML103 RILs, 159 CML333 RILs, 151 NC358 RILs, and 160 Tx303 RILs (Table S2). Table S3 lists the 413 lines used in the GWAS of leaf δ^13^C. Table S4 includes the QTL coordinates identified in the elemental QTL analyses. Figure S1 shows the distribution of leaf δ^13^C for each of the NAM RIL families. Figure S2 presents the correlation matrix for the elemental analysis, and Figure S3 shows the chromosomes where significant QTL were identified for each element. Figure S4 is the LOD plot from the GWAS mapping of leaf δ^13^C.

## RESULTS

### Single family QTL mapping

Previous studies investigating leaf δ^13^C in maize indicated that the NAM founder lines CML103, CML333, and Tx303 consistently contrast B73 with respect to leaf δ^13^C (Kolbe *et al.* 2018b; Twohey III *et al.* 2019). The founder line NC358 had a moderate leaf δ^13^C value (Kolbe *et al.* 2018b; Twohey III *et al.* 2019) and was also included in this study. The RIL families generated from these four founder lines were grown for linkage analysis. Consistent with previous studies, both the CML103 and CML333 parent lines had a significantly less negative leaf δ^13^C than B73 (*p* <0.05), when grown as replicated controls among the RILs. However, the Tx303 and NC358 parental lines were not found to be significantly different from B73. Transgressive segregation was observed in all four RIL families (Fig. S1).

Stepwise regression analyses found significant QTL for leaf δ^13^C in the NAM RIL families CML103, CML333, and Tx303 but not in NC358 (Fig. 1A). Interestingly, none of these QTL were shared between RIL families in this analysis. The CML103 RIL family had two significant QTL, one on chromosome 5 at 211.7 Mb and another on chromosome 7 at 142.4 Mb. Combined these two QTL accounted for 21.36% of the total phenotypic variance (Table 1). The RIL family CML333 had one significant QTL on chromosome 3 at 183.9 Mb, which accounted for 8.37% of the total phenotypic variation explained (Table 1). Finally, the Tx303 RIL family had a significant QTL on chromosome 2 at 13.5 Mb explaining 9.48% of phenotypic variation (Table 1). No significant QTL for leaf δ^13^C were identified in the NC358 RIL family.

**Table 1.** δ^13^C Single Family Stepwise Regression QTL

| Step | Marker | Family | Chr. | Position<br>(Mb) | *p*-value | TPVE*<br>(%) | Effect<br>Size | 1LOD Interval<br>(Mb) |
| --- | --- | --- | --- | --- | --- | --- | --- | --- |
| 1 | 824 | CML103 | 5 | 211.7 | 7.03E-06 | 12.32 | 0.1286 | 211.5 - 212.9 |
| 2 | 1032 | CML103 | 7 | 142.4 | 7.52E-05 | 21.36 | -0.1064 | 141.2 - 149.7 |
| 1 | 470 | CML333 | 3 | 183.9 | 2.07E-04 | 8.37 | 0.1137 | 178.6 - 195.7 |
| 1 | 251 | Tx303 | 2 | 13.5 | 6.04E-05 | 9.48 | -0.1457 | 13.5 - 15.2 |

\* Total Percent of Variation Explained

**Figure 1.**
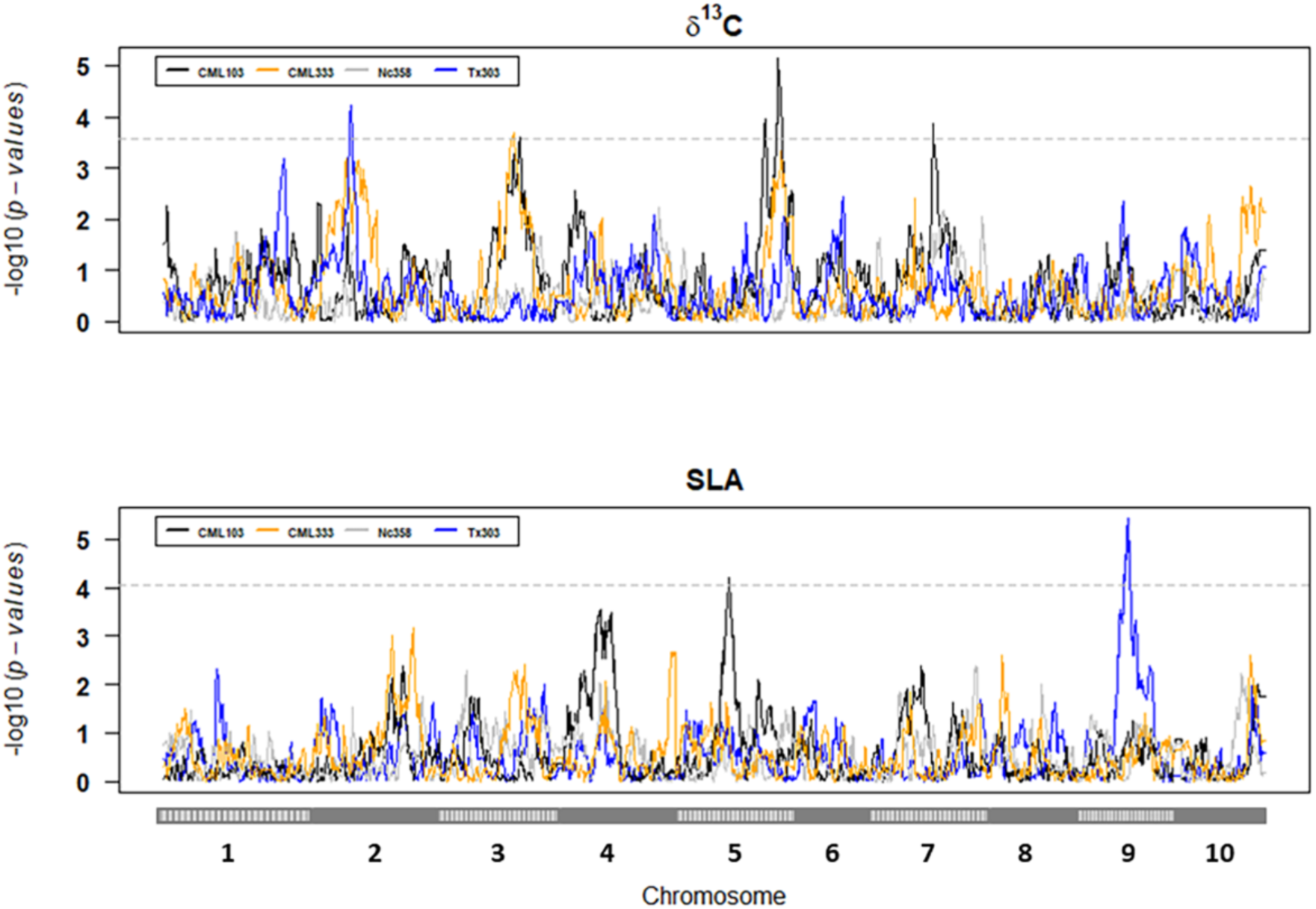
δ^13^C and SLA Single Family Stepwise Regression QTL Mapping. δ^13^C QTL (**A**) were identified in NAM RIL families CML103 (black), CML333 (orange), and Tx303 (grey) but not in NC358 (blue). Specific leaf area (SLA) QTL (**B**) were identified in NAM RIL families CML103 (black) and Tx303 (grey). Significance thresholds (dashed horizontal line) were determined by 200 permutations and an alpha of 0.05.

Specific leaf area was used as a proxy trait to test for a relationship between leaf thickness and leaf δ^13^C. No significant correlation was observed between SLA and leaf δ^13^C (*p* = 0.304). In addition to testing for a correlation with leaf δ^13^C, QTL mapping was performed for SLA to identify any possible overlaps with genomic regions identified for leaf δ^13^C. Mapping of SLA in the four RIL families identified two significant QTL. In the CML103 RIL family, a QTL was identified on chromosome 5 at 86.1 Mb and in the Tx303 RIL family a QTL on chromosome 9 at 107.8 Mb. Neither of the SLA QTL identified overlapped with QTL for leaf δ^13^C (Fig. 1B).

To test a potential link between transpiration and nutrient uptake, an elemental analysis was performed on leaf samples from each of the four RIL families. Samples were analyzed for 19 different elements using an IPC-MS. A full correlation matrix shows that some elements are highly correlated with each other (Fig. S2), but no strong correlations (*r* > +/- 0.7) were identified with δ^13^C. However, there was a weak but significant correlation (*p* = 6.745E-05, *r* = 0.18) between leaf δ^13^C and Mo (Fig. 2). Subsequent QTL mapping of the 19 element concentrations identified 28 QTL across 12 different elements (Fig. 3, Fig. S3). Significant QTL were found for B, Mg, P, S, K, Fe, Mn, Co, Cu, Rb, Sr, and Mo (Table S4). None of the elemental QTL overlapped with those found for leaf δ^13^C or SLA. However, in some cases multiple elements had common QTL, such as Mg and Mn on chromosome 10 in the CML103 RIL family and Co and Cu on chromosome 3 in the NC358 RIL family. Additionally, common QTL for an element were found across families, as in the case of Mg in the NC358 and Tx303 RIL families.

**Figure 2.**
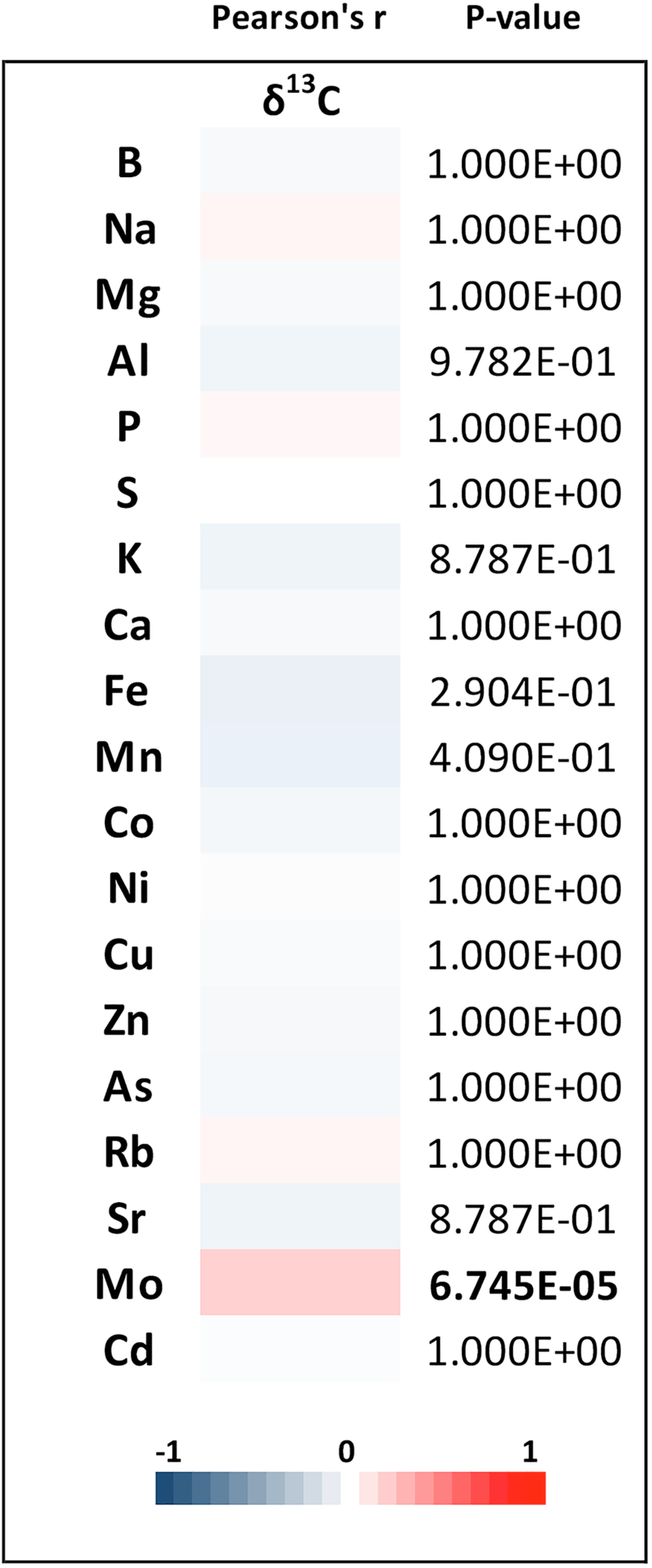
Pearson’s r Correlations. Correlations of mean phenotypic values using complete observations and Holm’s method to adjust *p*-values for multiple testing. The heat map shows no strong correlations between δ^13^C mean values and element mean values. δ^13^C and Mo have a significant p-value (p = 6.745E-05, r = 0.18).

**Figure 3.**
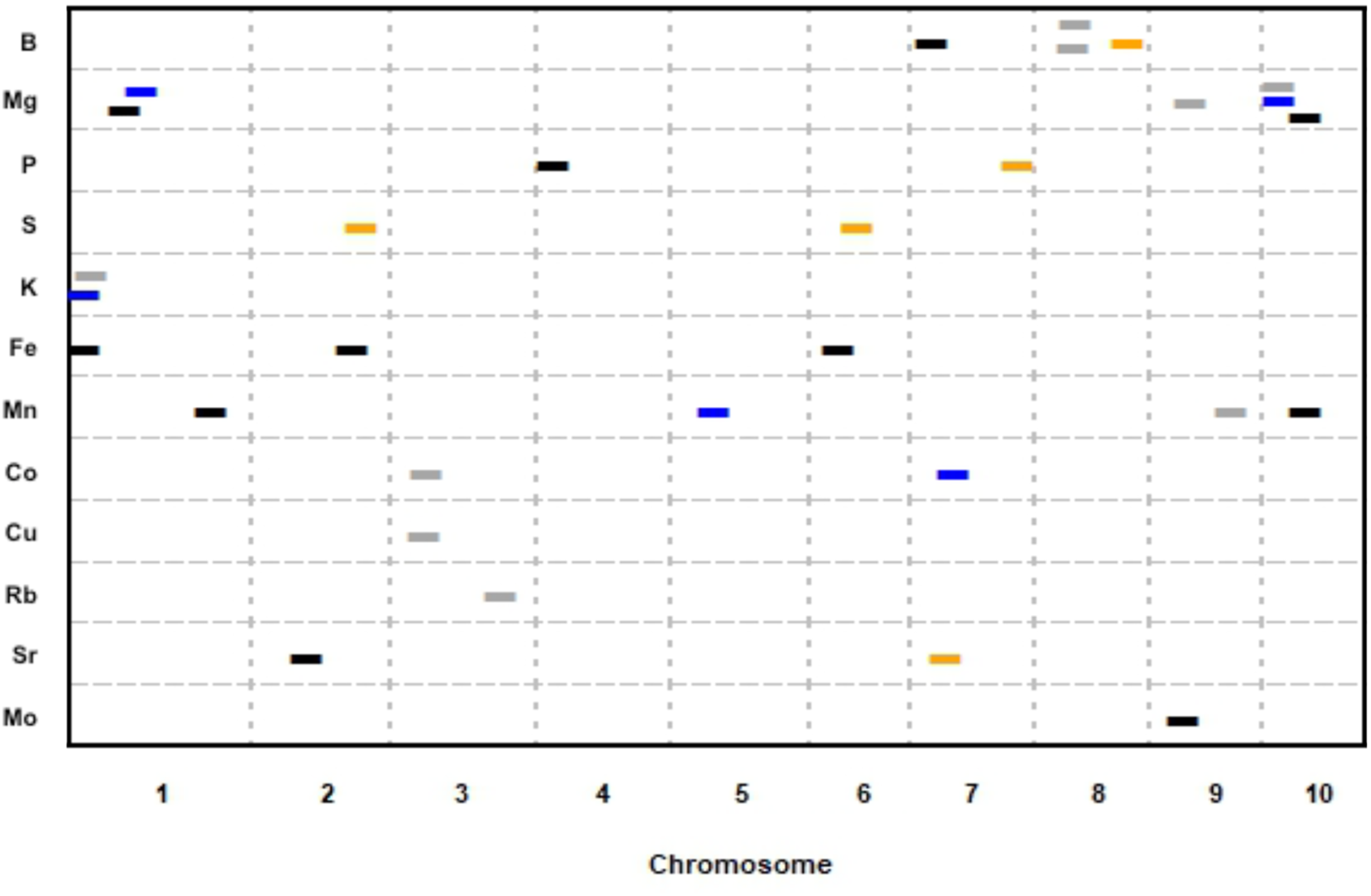
Element Single Family Stepwise Regression QTL Mapping. QTL mapping identified 28 QTL across 12 different elements. Significant QTL (alpha = 0.05) for each element are plotted. QTL location is shown across the 10 maize chromosomes (cM) on the x-axis. Dashes indicate a significant QTL, with the NAM RIL family in which the QTL was found designated by color; CML103 (black), CML333 (orange), Tx303 (grey), NC358 (blue). All dashes are the same length for visibility.

### Joint linkage QTL mapping

A joint linkage analysis was performed for leaf δ^13^C to test whether any additional QTL would be identified by combining the four RIL families into a single analysis. The joint linkage analysis identified the same significant QTL for leaf δ^13^C on chromosomes 2, 3, and 5 (Table 2) as in the single family stepwise regression analysis. Although the QTL on chromosome 7 was not found using the joint linkage approach, an additional significant QTL was identified on chromosome 1. Given that no significant QTL for leaf δ^13^C were identified in the NC358 RIL family, we tested whether removing this family from the joint linkage analysis would change the outcome. When the joint linkage analysis was rerun excluding family NC358, the same four QTL were reidentified with decreased *p*-values, and the total phenotypic variation explained (*R*^2^ value) increased in later steps of the model. However, no new QTL were identified with this approach.

**Table 2.** δ^13^C Joint Linkage Mapping QTL

| All Four Families |  |  |  |  |  |  |  |  |
| --- | --- | --- | --- | --- | --- | --- | --- | --- |
| Step | QTL | Family | Chr. | Position<br>(Mb) | *p*-value | TPVE*<br>(%) | Effect<br>Size | 1LOD Interval<br>(Mb) |
| 1 | m0200 | Tx303 | 2 | 13.8 | 2.82E-06 | 7.9 | -0.144 | 12.6 - 15.8 |
| 2 | m0677 | CML103 | 5 | 211.2 | 1.98E-04 | 11.8 | 0.123 | 211.2 - 212.7 |
| 3 | m0385 | CML103 | 3 | 182.1 | 1.44E-04 | 15.24 | 0.115 | 180.0 - 195.3 |
| 4 | m0132 | Tx303 | 1 | 263.2 | 4.41E-05 | 18.19 | -0.126 | 257.1 - 263.6 |

| Excluding NC358 |  |  |  |  |  |  |  |  |
| --- | --- | --- | --- | --- | --- | --- | --- | --- |
| Step | QTL | Family | Chr. | Position<br>(Mb) | *p*-value | TPVE*<br>(%) | Effect<br>Size | 1LOD Interval<br>(Mb) |
| 1 | m0200 | Tx303 | 2 | 13.8 | 5.70E-06 | 7.47 | -0.144 | 12.6 - 15.9 |
| 2 | m0677 | CML103 | 5 | 211.2 | 2.86E-04 | 12.31 | 0.123 | 211.2 - 212.7 |
| 3 | m0385 | CML103 | 3 | 182.1 | 1.60E-04 | 16.45 | 0.115 | 180.0 - 195.3 |
| 4 | m0132 | Tx303 | 1 | 263.2 | 5.97E-05 | 20.05 | -0.121 | 253.0 - 263.6 |

### Genome wide association study

Once significant QTL intervals were identified for leaf δ^13^C using a biparental mapping strategy (Fig. 1), we performed a genome wide association study to try and narrow down the intervals to specific genic regions. The Wisconsin Diversity Panel was chosen because it represents a large portion of variation found within maize and has a robust publicly available 485,179 SNP set. A subset of 413 of the possible lines were chosen due to seed availability, and were grown in a single randomized block. No significant SNP associations with leaf δ^13^C were identified (Fig. S4).

## DISCUSSION

Leaf δ^13^C has a moderately high heritability in maize (Twohey III *et al.* 2019), which facilitates the use of quantitative genetics approaches to pinpoint the genomic locations controlling this trait. Here we characterized the genetic control of δ^13^C in maize using leaf tissue collected at vegetative stage V9-V10 to reflect the photosynthetic pool during active growth. We were able to identify several significant QTL for leaf δ^13^C across three NAM RIL families using stepwise regression. Using these populations we were also successful in identifying QTL for SLA and 12 different elements. Contrary to our hypothesis, no significant correlation was observed between leaf δ^13^C and SLA or elemental composition.

We strategically picked NAM RIL families based on the founder parents that had the largest differences in their δ^13^C for single family and joint linkage analyses. However, we also included a parent which was not extremely different from B73. Interestingly, we were unable to identify significant QTL in the NC358 RIL family despite transgressive segregation. We interpret this result as an indication the NC358 contains only small effect QTL that were not detected in this study. Alternatively, NC358 leaf δ^13^C may be more sensitive to the growing environment with a smaller genetic component. Twohey III *et al.* noted that while several maize lines were stable when tested in greenhouse and field environments, there were other lines that had highly variable isotopic signatures (2019). A large amount of environmental influence over this trait in some backgrounds would obscure the genetic contribution and our ability to detect significant QTL.

When we compared the regions identified here with regions previously mapped in *S. viridis* no obvious overlap was observed (Ellsworth *et al.*, 2020). However, of the QTL identified for kernel δ^13^C in maize (Gresset *et al.* 2014, Avramova *et al* 2019), our QTL for leaf δ^13^C overlapped those on chromosomes 1, 3, and 7. This result demonstrates that some QTL for δ^13^C are shared between tissues, and that these QTL are identified across several populations and environments. Therefore, while metabolic processes have the possibility of influencing the δ^13^C as products are mobilized from source to sink tissues, our data would suggest that the initial source signature is maintained to a large degree in the kernel.

Although the QTL analyses presented here do not provide gene-level resolution, we were able to look for candidate genes within the intervals. The chromosome 5 QTL includes an Erecta-like gene (*er1*, GRMZM2G463904, 211.8 Mb). Unfortunately, stomatal density data were not collected on these populations, which would further support the role of *er1* in variation of leaf δ^13^C. This would be an interesting area of future research given that this gene was found to effect δ^13^C in Arabidopsis by changing stomatal density (Masle *et al.* 2005). We also looked for genes that have been previously shown to directly influence transpiration efficiency in maize (reviewed in Leakey 2019). However, none of these were found to be located in our QTL intervals.

The linkage analyses using biparental mapping populations identified several significant QTL, but none of the single family QTL were independently identified in more than one family (Fig. 1). This result indicates that leaf δ^13^C can be controlled by different factors depending on the genetic background. Furthermore, if in fact leaf δ^13^C is controlled by many small effect QTL, this may explain why the GWAS did not identify any significant SNP associations with leaf δ^13^C. Identifying rare alleles with small to moderate effect size is a known weakness of the GWAS method (Bazakos *et al.* 2017). A better understanding of the mechanisms influencing leaf δ^13^C would allow future analyses to move beyond single marker tests and instead look at SNPs in genes representing a particular pathway or process that could be collectively significant. This approach was successfully used to study maize lipid biosynthesis (Li *et al.* 2019).

The diffusion of CO_2_ into mesophyll cells is a potential source of variation in leaf δ^13^C, which could be linked to stomatal density or leaf thickness. Previous work in maize has shown that stomatal density is not correlated with leaf δ^13^C in a small diversity panel of maize (Foley 2012). SLA has not been linked to δ^13^C in maize, but in rice δ^13^C and SLA have shared QTL (This *et al.* 2010). In this study we were able to test SLA and δ^13^C in four RIL families, and no correlation was observed. Likewise, a comparison of the QTL analyses showed no overlapping regions for SLA and those mapped for leaf δ^13^C. This result suggests two possibilities. First, it is possible that differences in SLA observed in these populations are not due to leaf thickness, but rather composition. Identifying the causative genes underlying the QTL would give insight into the mechanism. A second possibility is that leaf anatomical traits other than leaf thickness influence leaf δ^13^C. A variety of anatomical traits could affect δ^13^C and would not be captured by measurement of SLA.

With our data we were able to indirectly test the relationship between nutrient uptake and transpiration. If reducing transpiration limits nutrient uptake, transpiration efficiency as a trait for increasing WUE would have limited application. The nineteen elements tested here were previously reported to have narrow-sense heritabilities ranging from 0.11 to 0.66 (Baxter 2014). The only element found to be significantly correlated to leaf δ^13^C was Molybdenum. Molybdenum is required for several vital biological processes related to nitrogen and water (Baxter 2008). Because molybdenum is a required cofactor for ABA synthesis, maize plants overexpressing molybdenum cofactor sulfurase gene have increased drought tolerance (Lu *et al.* 2013). However, in our study we observed a positive correlation between leaf δ^13^C and molybdenum, which is contrary to expectation given that an increase in δ^13^C signifies a decrease in WUE. Overall, it is encouraging that the majority of elements sampled were not associated with δ^13^C. This suggests that breeding for leaf δ^13^C as a means to reduce transpiration would be unlikely to result in plants with nutrient uptake deficiencies.

Although the main focus of this study was to investigate leaf δ^13^C and its relationship to SLA and nutrient accumulation, the QTL mapping of the analyzed elements was an interesting biproduct. Mapping the leaf ionome of the four RIL families resulted in many significant QTL, including some overlapping intervals for different elements. Multi element QTLs are common, and are thought to be due to loci affecting processes such as the acidification of the rhizosphere or altering the permeability of the casparian strip (Baxter 2015). Interestingly there was not much overlap between the ionomic QTL identified here and a previous study on kernels (Table 4 Baxter *et al.* 2014). The only overlapping QTL was for Rb85 on chromosome 3. There are several possible causes for the limited overlap between these methods. The ionome is strongly influenced by geneotype by environment interactions, with many of the QTL identified in previous studies being location specific (Asaro *et al.* 2016). Additionally, there could be differences between the leaf and grain ionome is due to differential mobilization of nutrients from vegetative tissues into kernels during grain fill.

## Acknowledgements

We thank Aaron Slack, Robert Twohey III, and Mengqiao Han for technical assistance with growing and sampling the mapping populations. This work was supported by United States Department of Agriculture - Hatch, a United States Department of Agriculture - Agriculture and Food Research Initiative grant (2019-67013-29195), the United States Department of Agriculture - Agricultural Research Service (5070-21000-039-00D), as well as a Department of Energy, Office of Science, Office of Biological and Environmental Research grant (DE-SC0018277).

---

**Supplemental Figure 1.**
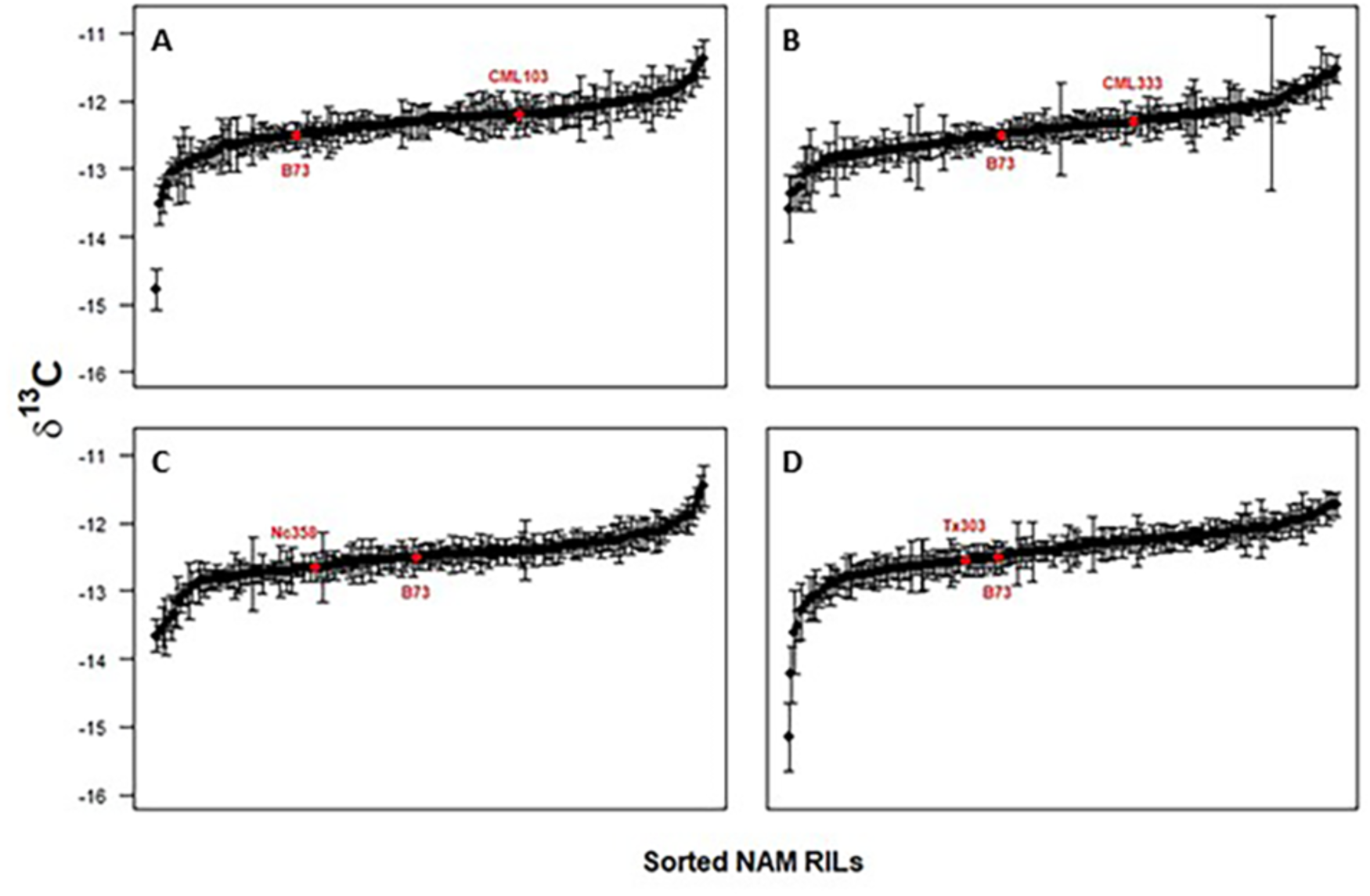
NAM RIL Transgressive Segregation. NAM RIL families CML103 (**A**), CML333 (**B**), NC358 (**C**), and Tx303 (**D**) were sorted by δ^13^C and plotted. Parental lines are shown in red.

**Supplemental Figure 2.**
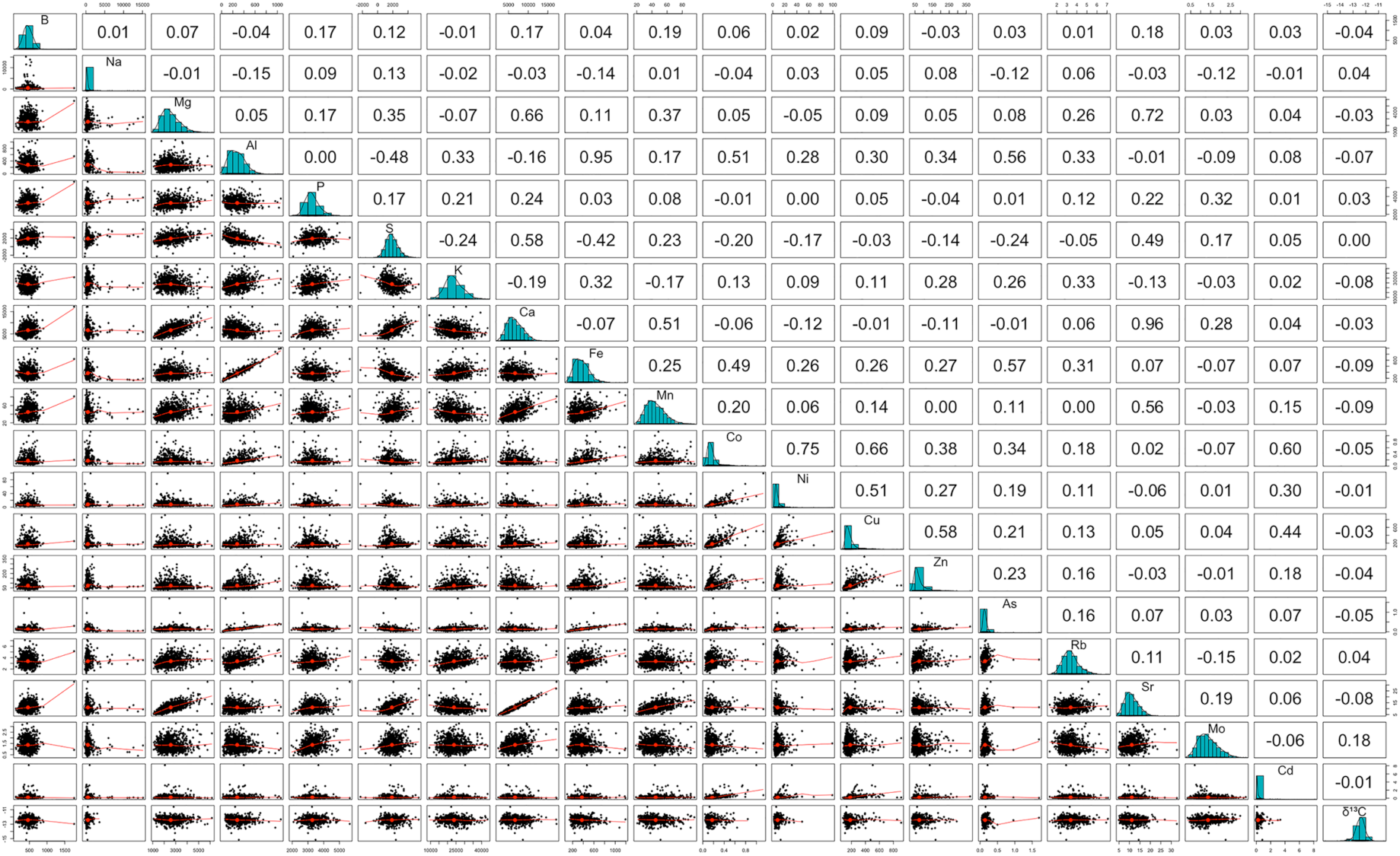
Element and δ^13^C Full Correlation Matrix. A full correlation matrix of element and δ^13^C mean values is show. The diagonal displays histograms of each dataset. Pearson’s r is shown in the upper panel. Scatter plots and best fit line are shown in the lower panel.

**Supplemental Figure 3.**
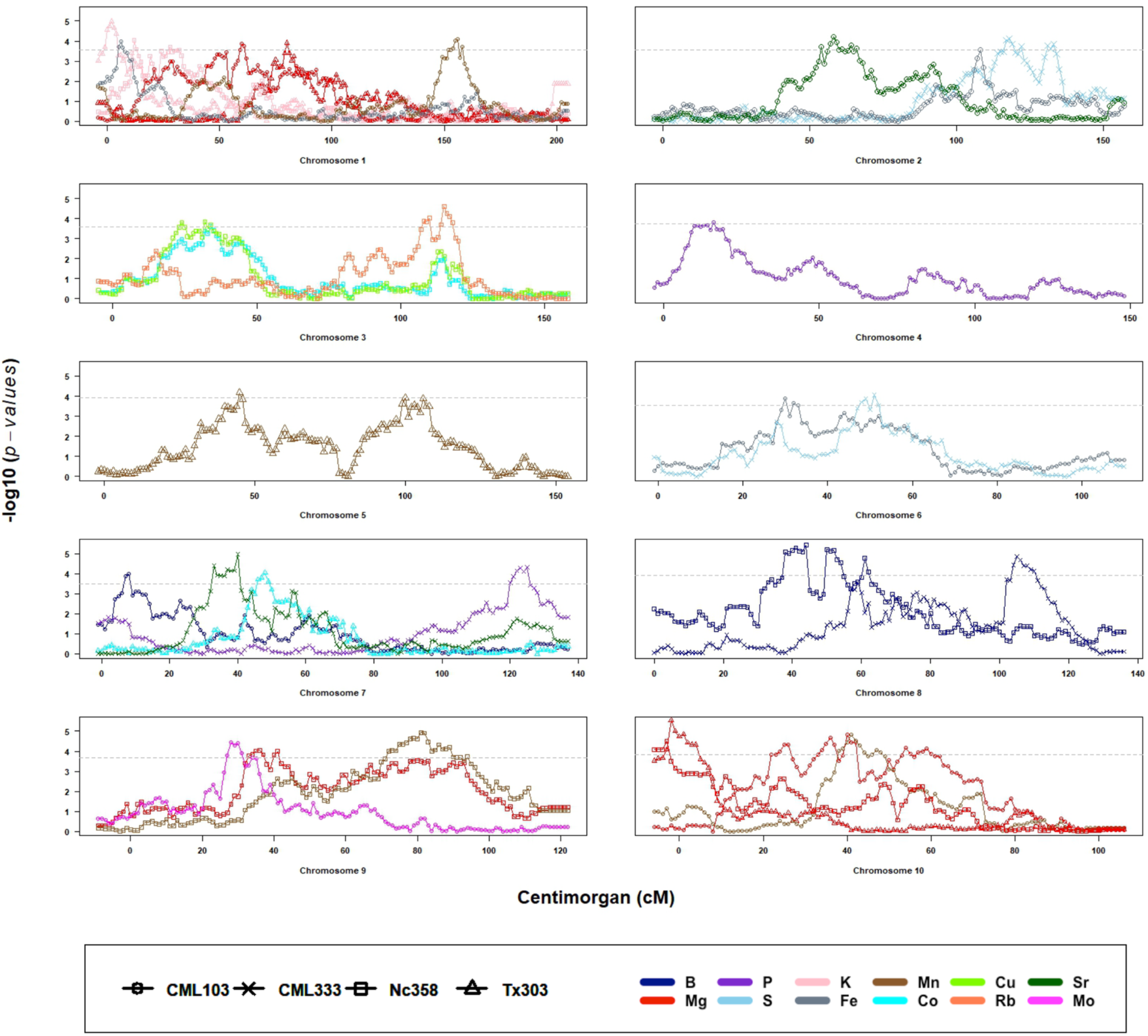
Element QTL Mapping by Chromosome. Significant element QTL are shown by maize chromosomes 1 through 10 on the x-axis (in cM). Each NAM RIL family is represented by a symbol; CML103 (○), CML333 (x), NC358 (□), and Tx303 (Δ). Each element is designated by color. Significance thresholds (dashed horizontal line) were determined using 200 permutations, alpha=0.05 for each QTL independently.

**Supplemental Figure 4.**
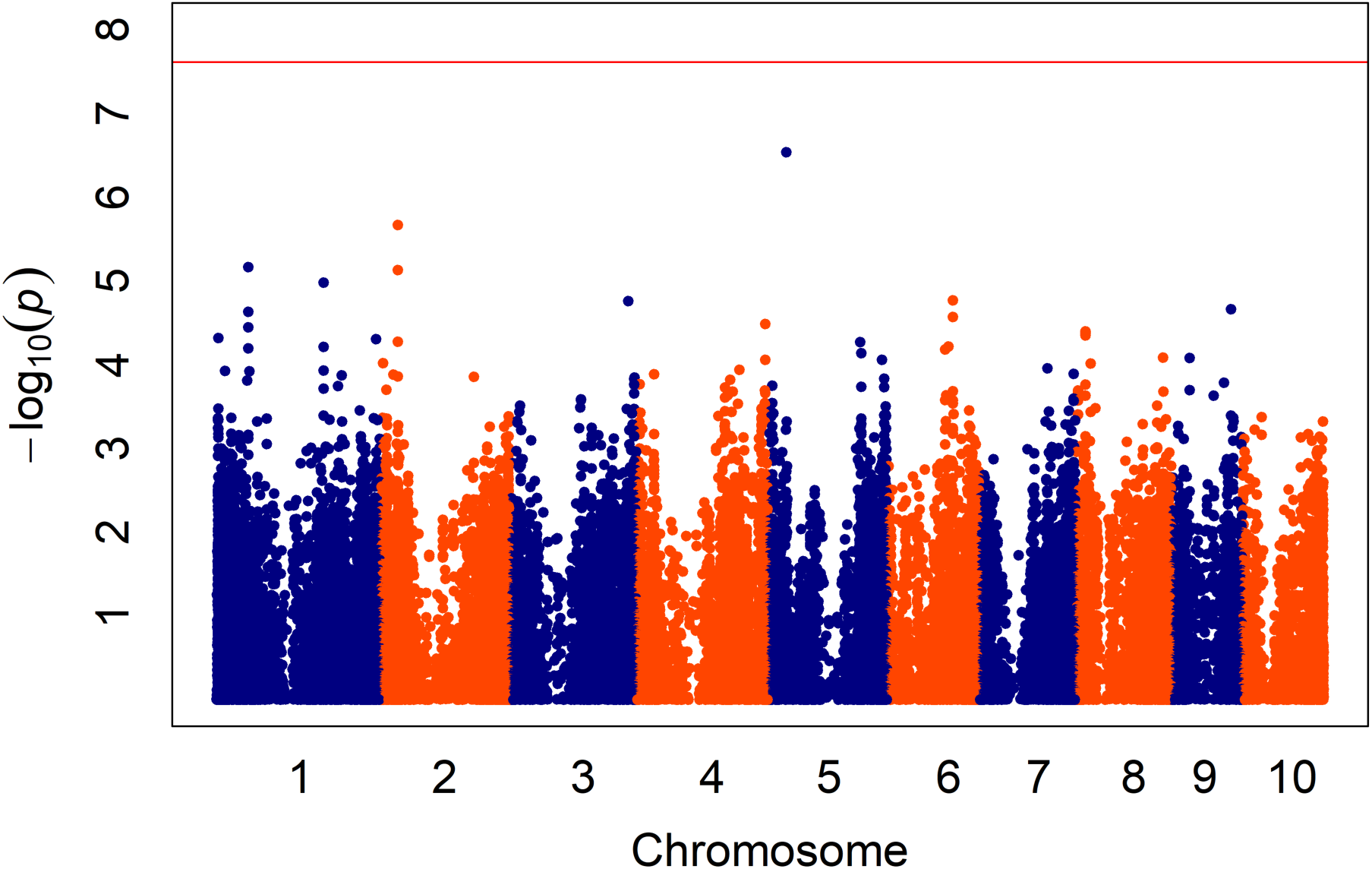
Genome Wide Association Study for δ^13^C. Manhattan plot showing significance of SNPs derived from a mixed linear model using the Bayesian information criterion to select the optimal number of principal components. The significance threshold represents the Bonferroni correction of familywise error rate.

